# Connecting adhesion dynamics and trail formation in malaria parasites by imaging the major surface antigens CSP and TRAP

**DOI:** 10.64898/2026.08.26.747238

**Authors:** Kevin Walz, Mirko Singer, Leon Lettermann, Yvonne Sokolowski-Adams, Carolina Thieleke-Matos, Sylvia Olberg, Magdalena C. Unterreiner, Christine Selhuber-Unkel, Vibor Laketa, Ulrich S. Schwarz, Friedrich Frischknecht

## Abstract

Malaria infections are initiated by mosquito bites, during which *Plasmodium* sporozoites are injected into the host skin. Sporozoites migrate rapidly to find and enter blood capillaries and ultimately invade hepatocytes. Sporozoite migration and invasion is mediated by the transmembrane protein thrombospondin-related anonymous protein (TRAP), which links the extracellular substrate to the actomyosin complex powering gliding motility, while the abundant, GPI-anchored circumsporozoite protein (CSP) covers most of the parasite membrane and modulates adhesion. Both proteins are secreted onto the parasite surface and deposited in a membranous trail originating at the parasite rear. The surface dynamics of these essential sporozoite proteins and the mechanism of deposition, however, are not understood. Here, using orbital total internal reflection fluorescence microscopy (TIRF), we reveal the dynamics of TRAP adhesion site formation and disassembly as well as CSP and TRAP deposition rates. We find that TRAP assembles into distinct adhesion sites, which then undergo retrograde translocation as the sporozoite moves forward. Around half of the TRAP adhesins, together with CSP, remain associated in small membrane droplets on the substrate after the sporozoite has disengaged from the adhesion site. These droplets seem to originate from nanotubes, that presumably decay under high tension. Strikingly, we observe a change in actin filament accumulation if proteolytic cleavage of TRAP is inhibited, providing the first visual evidence for outside-in signaling in sporozoites. Our study reveals a relation between adhesion dynamics and trail formation in *Plasmodium* sporozoites that might also be relevant for other cell types.

## Introduction

Gliding motility is essential for apicomplexan parasites to quickly migrate on and through tissues as well as for host cell invasion. Contrary to crawling motility, gliding does not involve a change of cell shape and often exceeds speeds of one micrometer per second. The machinery facilitating gliding motility is termed the glideosome and centers around relatively short actin filaments propelled backward by myosin A (myoA), a single-headed myosin motor. The glideosome resides just beneath the plasma membrane and is anchored in the inner membrane complex (IMC), a flattened vesicular organelle that subtends the plasma membrane of the alveolate at a distance of around 30 nm (*1, 2*). The actin filaments are likely linked to extracellular ligands through membrane-spanning adhesins, such as those of the TRAP family (*3, 4*). Adhesins are stored in micronemes, secretory vesicles that release their content at the apical pole of the parasite (*5–8*). The forward motion in gliding motility is generated through the apical secretion of adhesins and their actin-mediated rearward flow after binding to extracellular substrates (*1, 9–11*). For motility of *Plasmodium* sporozoites, the thrombospondin related anonymous protein (TRAP) is essential as well as for salivary gland and liver invasion (*12, 13*). Substrate adhesion is speculated to take place in distinct adhesion sites initiated with the secretion of a single microneme. However, TRAP containing adhesion sites have not yet been directly visualized. For *Toxoplasma gondii* tachyzoites moving through matrigel, circular constrictions zones were shown, potentially suggesting circular adhesion sites during 3D motility (*14*). Using micropatterning of adhesive and non-adhesive substrates, it was shown that *T. gondii* tachyzoites can generate forward motility in 2D with a single adhesion site (*15*). During 2D motility, *Plasmodium* sporozoites are also believed to move over individual adhesion sites, as was suggested previously with reflection interference contrast microscopy (RICM) (*13*). Curiously, only very few adhesins are present at the surfaces of parasites, which are dominated by GPI-anchored proteins, such as surface antigen 1 (SAG1) on *T. gondii* tachyzoites, merozoite surface protein 1 (MSP1) on *Plasmodium* merozoites and circumsporozoite protein (CSP) on *Plasmodium* sporozoites (*16–18*). CSP is essential for sporozoite formation in oocysts at the mosquito midgut, egress from oocysts, entry into salivary glands, migration within the skin and entry into hepatocytes (*19–24*). CSP is also the basis of the two licensed malaria vaccines and anti-CSP antibodies inhibit migration in the skin, and thus ultimately preventing liver invasion (*25–29*).

TRAP contains two adhesive extracellular domains, a transmembrane domain and a cytoplasmic tail that links to the actin cytoskeleton (*30, 31*). TRAP is cleaved by a rhomboid protease, likely ROM4, as the *T. gondii* homolog of TRAP, MIC2, is cleaved by ROM4 (*32, 33*). This cleavage is essential for efficient migration and TRAP accumulates on the surface in the absence of cleavage (*32*). CSP consists of an unstructured N-terminal domain, a central repeat region and a C-terminal adhesion domain that is in turn linked to a GPI anchor for integration into the plasma membrane (*34*). Over the course of sporozoite progression in the mosquito and specifically during liver invasion, the N-terminus is cleaved at a short proteolytic cleavage site termed region I by an unknown protease (*21, 22*). CSP is also found in trails left behind by moving sporozoites, a phenomenon well established to visualize gliding capabilities of sporozoites (*32, 35*). Similar, SAG1-containing trails are also found behind gliding *T. gondii* tachyzoites (*36*). Trail formation is also very common for crawling cells, which often show a broad region of membrane nanotubes (sometimes called retraction fibers) being pulled out of their rear, where mature adhesion sites and strong actomyosin contractility pin the cell to the substrate (*37–39*). Like for malaria parasites, proteolytic cleavage is also an important aspect of rear detachment and mediated by proteases such as calpain, matrix metalloproteinases and sheddases of the ADAM family. In *Plasmodium* sporozoites with their slender shapes, similar processes are strongly localized to one point, making them an interesting model system to study trail formation.

How TRAP forms adhesion sites and links to actin filaments, how CSP behaves in the sporozoite membrane and how both are linked to trail formation at the rear are open questions that could be answered using fluorescently tagged proteins. Both TRAP and CSP were studied extensively with antibodies (*22, 32, 40*) and parasite lines expressing GFP-tagged versions have been generated. While endogenous TRAP could be GFP-tagged between the signal peptide and the first adhesion domain without a loss of infectivity (*41*), tagging of CSP proved more difficult and did not lead to mature sporozoites that could be imaged (*42, 43*). Similarly, the visualization of actin filaments proved difficult with GFP-tagged actin only providing limited insights (*44*). Visualization of actin filament accumulations was made possible only with the generation of parasite lines expressing an actin filament-recognizing nanobody linked to a fluorescent protein (*45, 46*).

Here we address the open question how adhesion dynamics and trail formation are related in gliding *Plasmodium* sporozoites. Currently, trail formation is believed to be caused by protein shedding (*47*), commonly interpreted as a direct deposition of protein as a result of proteolytic cleavage, antibody mediated crosslinking or potentially cleavage by a GPI anchor specific phospholipase. To investigate trail formation in spatiotemporal detail, we successfully GFP-tagged CSP in a clonal parasite population that allows imaging of mature sporozoites. To visualize surface proteins dynamically we established orbital ring total internal reflection fluorescence (oTIRF) microscopy. This microscopy modality provides an evenly illuminated evanescent field for sensitive surface imaging providing the necessary contrast and resolution (*48*). oTIRF allowed us to quantitatively image the formation and dynamics of TRAP-containing adhesion sites showing that each adhesion site contains multiple TRAP molecules. The deposition of CSP and TRAP in trails during sporozoite motility results from incompletely disassembled adhesion sites that form membrane nanotubes at the sporozoite rear, that then decay into droplets and form the trail.

## Results

### CSP deposition in trails can be visualized by TIRFM

While sporozoites move in right-handed helical paths in three-dimensional environments, their movement on a two-dimensional surface in medium occurs in counter-clockwise circles, likely because adhesins are preferably secreted at the titled apical ring towards the side that forms stable attachment with the substrate (*10, 49*). Reflection interference contrast microscopy revealed that parasite migration is mediated through dynamic formation and turn-over of distinct adhesion sites (*13, 24*) (Figure 1A). These adhesion sites are likely composed of TRAP protein family members embedded in a sea of CSP proteins and linking the extracellular substrate to the actomyosin complex (Figure 1B). To visualize CSP during motility of salivary gland sporozoites, we generated a parasite line expressing an additional copy of CSP, internally fused to GFP (Figure 1C). To avoid previous detrimental phenotypes caused by tagging of CSP (*42, 43*), the additional copy was integrated into a silent genomic region on chromosome 12 (*50*) (Supplemental Figure 1). This copy was placed under the *csp* promoter, but featured a shortened 3’UTR to limit expression of the fluorescent protein (*43*). The resulting CSP-GFP parasite line showed similar numbers of sporozoites in oocysts and salivary glands to wild type parasites (Figure 1D). Sporozoites of the line migrated in similar fashion and at similar speed to wild type parasites (Figure 1E,F). Western Blot analysis shows two additional GFP-containing bands in CSP-GFP salivary gland sporozoites compared to the two wild type bands, likely corresponding to the additional Region I-processed and uncleaved CSP-GFP versions (Figure 1G). Previously, deposition of CSP containing trails was only shown for motility on two-dimensional surfaces and it was therefore unknown whether this phenomenon would also occur during motility *in vivo*. To visualize possible trail formation in a 3D environment, more similar to *in vivo* than the typical 2D *in vitro* assays, we performed immunofluorescence with α-CSP antibodies in isotropic polyacrylamide gels after migration of sporozoites (*10, 51*). This revealed spiral patterns similar to the helical paths of sporozoites moving in 3D (Figure 1H, Supplemental Figure 2). To visualize the live deposition of CSP-containing trails, a strong signal-to-background (and trail to sporozoite) ratio is required. Regular epifluorescence imaging of CSP-GFP expressing sporozoites only allows the visualization of strong trails. The selective excitation of surface-close fluorophores in oTIRF microscopy also reveals CSP-GFP containing trails of lower intensity (Figure 1I) as the majority of CSP-GFP on the surface of the sporozoite is not exited, allowing for live imaging of CSP-GFP trail deposition in gliding sporozoites. We therefore continued the analysis of trail formation using this oTIRF 2D *in vitro* assay.

**Fig. 1:**
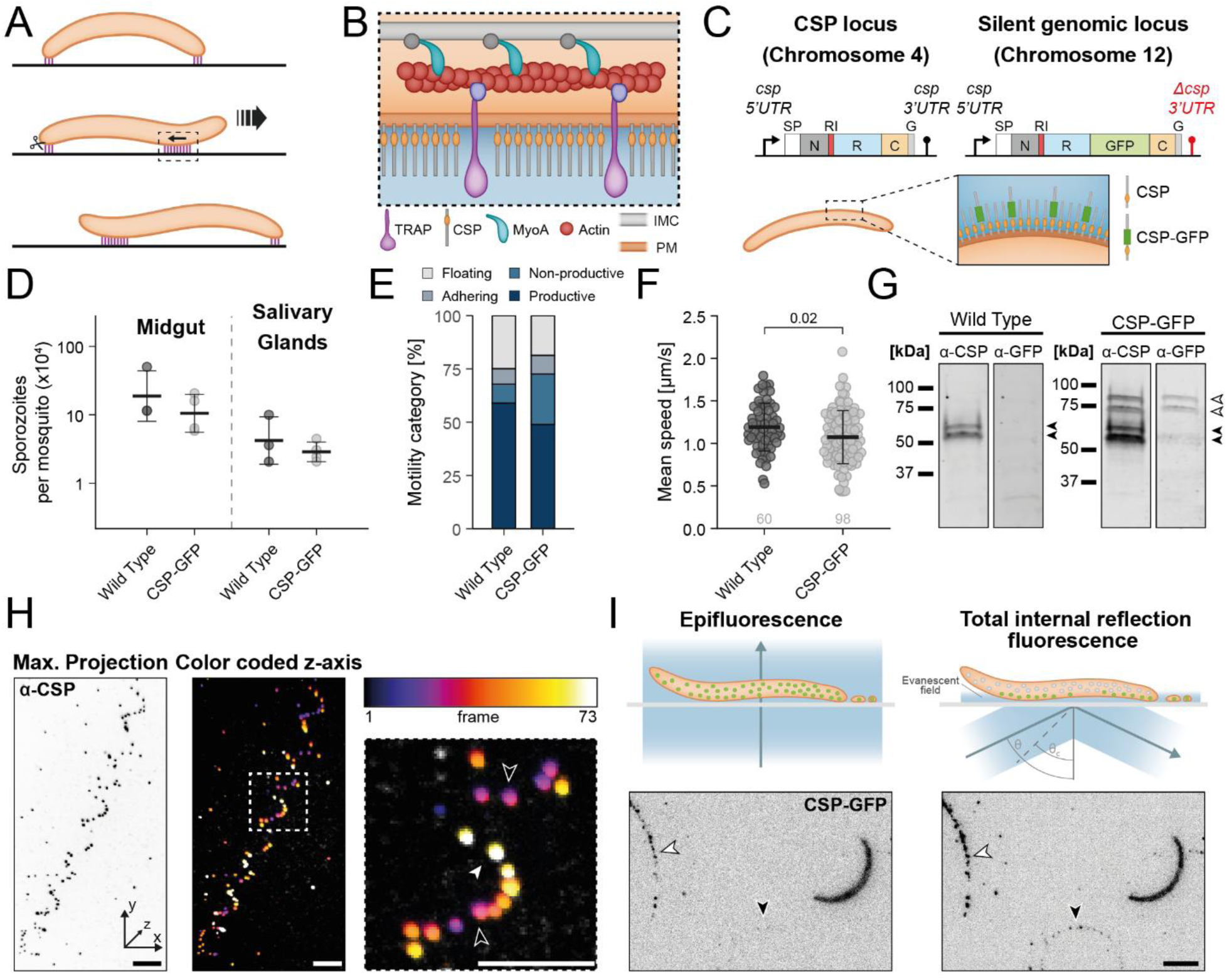
Fluorescent tagging of CSP allows for live visualization of trail formation. **(A)** Sporozoites gliding motility through stick-and-slip motility requires dynamic turn-over of adhesion sites. Rearward moving adhesion sites are bound to surface ligands, generating forward motion. Posterior sites are disassembled to continue motility. **(B)** Simplified structure of the glideosome at an adhesion site consisting of inner membrane complex (IMC)-bound myosin A (MyoA) motors moving actin filaments that link to plasma membrane (PM)-spanning TRAP adhesins surrounded by CSP molecules. **(C)** Strategy for expression of an additional copy of CSP-GFP, integrated into a silent genomic region on chromosome 12, utilizing a shortened CSP 3’ UTR (*Δcsp 3’UTR*). Schematic illustrating the estimated ratio of CSP and CSP-GFP molecules on the parasite surface. **(D)** Number of sporozoites in the midgut and salivary glands of infected mosquitos. Each data point represents a separate infected cage. Number of analyzed mosquitos: Wild type = 58; CSP-GFP = 79. **(E)** Categories of motility patterns present in gliding sporozoites. Data consists of two pooled separate replicates. Number of analyzed sporozoites: Wild type = 904; CSP-GFP = 1186. **(F)** Mean speed of productively gliding sporozoites. Data consists of two pooled separate replicates. Significance is calculated through pair-wise comparison with Wild Type in Student’s t-test. The p-value is shown directly. **(G)** Western Blot analysis of salivary gland sporozoites. Samples were probed using a α-CSP repeat antibody and an α-GFP antibody. The expected double bands of the native CSP (black arrowheads) and of the CSP-GFP (white arrowheads) are marked. **(H)** Immunofluorescence of CSP in hydrogel environment after sporozoite motility using a spinning disc microscope. Illustrative helical trail of CSP signal is shown as z-projection and depth-colored projection. Scale bar: 10 µm. I) Schematic illustration of selective excitation of fluorophores in total internal reflection fluorescence (TIRF) microscopy as well as exemplary images of a CSP-GFP expressing sporozoite and trails. The position of trails with strong signal (white arrowhead) and weaker signal (black arrowhead) are marked. Scale bar: 5 µm.

Dynamic imaging of CSP-GFP expressing sporozoites by oTIRF reveals the process of trail formation to be mediated by the extrusion, stretching and collapse of CSP-GFP containing, membrane tube-like structures (Figure 2A-C, movies 1-3). The formation of nanotubes is a very common process in cells and is often combined with membrane theory to measure the material properties of membranes, in particular bending stifffness k and surface tension s. With typical values k=25 k_B_T and s=0.1 mN/m, the tube diameter should be 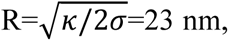 which is well below the resolution limit (*52, 53*). These tubes can stretch over relatively long distances, before being separated from the sporozoite and collapsing even further into smaller trail fragments (Figure 2A, movie 1). This collapse is typical for membrane tubes under tension and is an example of the Rayleigh-Plateau instability of cylinders under tension (*54*). Typically, the radius of the droplets should be twice the radius of the tubes, that is around 46 nm. Trails can also be seen in scanning electron microscopy (SEM) and SEM indeed revealed similar vesicular droplets (Supplemental Figure 3). The position of the trail frequently also seems to remain static even if parts of a tube collapse further, indicating some stable anchoring of the trails to the substrate. The most abundant type of trail formation consists of small bleb-like extrusions being pulled away from the moving sporozoite and being placed as circular trail fragments after the collapse of the connection (Figure 2B, movie 2). In a few sporozoites that do not productively move, but rather only move in short back-and-forth motion, the trails can stay attached to the sporozoite and even form on the rear and front (Figure 2C, movie 3). While these tethers could possibly stall sporozoite migration, pulling them backwards, they appear like passive connections to the surface that remain attached as the sporozoite cannot move forward. This elastic behavior suggests that these trails consist of the remnants of collapsed membrane tubes.

**Fig. 2:**
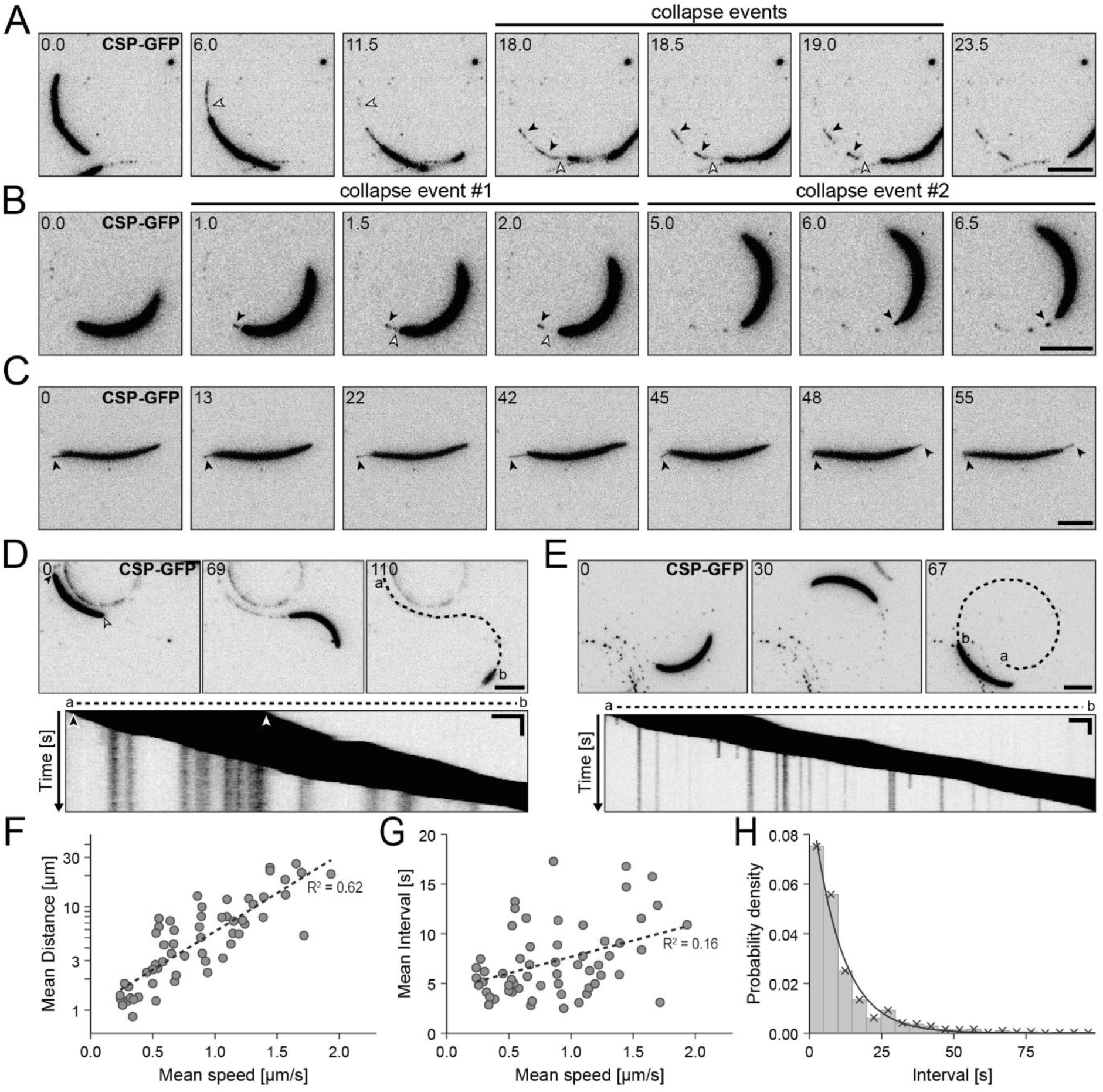
Visualizing CSP-GFP in trail formation reveals the process to consist of stretching and collapse of membrane protrusions. **(A-C)** Time lapse of gliding sporozoites expressing CSP-GFP in oTIRF with **(A)** an extreme example of a sporozoite displaying long extrusions of membrane that collapse into multiple trail depositions, **(B)** an exemplary sporozoite forming multiple subsequent extrusions that collapse into single deposits and **(C)** a patch gliding sporozoite with stable tethering connections on both ends extending and shrinking during the forward and back motion. The position of anchored deposits (black arrowhead) and collapsing tethers (white arrowheads) are marked. Scale bar: 5 µm. **(D-E)** Generation of a kymograph for trail analysis on two exemplary sporozoites. The sporozoite path is tracked (dashed line) from the start (a) to the end (b) of the path. The resulting kymograph consists of the intensity along the path for each time point. The position front (white arrowhead) and back (black arrowhead) of the sporozoite in the first frame are marked in the image and kymograph respectively. Scale bar: 5 µm. **(F)** Mean distance between trail deposits of 58 sporozoites versus their mean speed and linear regression in the logarithmic space. **(G)** Mean distance between trail deposition events of 58 sporozoites versus their mean speed and linear regression (dashed line) with the R^2^. **(H)** Histogram of the interval between the deposition events of 58 sporozoites and an exponential fit with the R^2^.

### Trail deposition is a stochastic process

To quantitatively analyze the trail formation process, we utilized kymographs. To this end, we measured the fluorescence signal along the path of the moving sporozoite and plotted the intensity over time (Figure 2D,E, movies 4,5). This visualization allows for the identification of deposited trail fragments as vertical lines extending beyond the diagonal signal of the moving sporozoite (Figure 2D, movie 4). In some scenarios, the trail fragments also display sudden dissociation from the surface, suggesting their connection to the substrate is lost (Figure 2E, movie 5). Correlating these trail characteristics to the gliding capabilities of the sporozoite may reveal the connection between trails and the gliding machinery. Plotting the mean distance between newly established trail fragments against the mean speed of 58 migrating sporozoites showed a clear positive correlation (Figure 2F). However, there was no correlation between sporozoite speed and the time interval of the moment of trail fragment formation (Figure 2G). This suggests that less material per length is deposited at higher speeds, but that the droplet sizes are similar, possibly because they result from a Rayleigh-Plateau instability with a critical wavelength that is independent of speed.

The deposition of membrane trails may also be a function of accumulating excess plasma membrane, for example, through the secretion of micronemes. To maintain an equilibrium, this excess membrane may be left behind once a critical amount of it accumulates on the surface. In this scenario we would expect the rate of trail formation to behave in a time-dependent matter. We examine whether intervals between trail deposition events behave stochastic or predominantly in a time-dependent way, which would be expected for accumulation of excess membrane. A regular time-dependent mechanism would produce a narrowly distributed, highly repeatable spacing. Instead, we observed substantial variability, with a coefficient of variation CV = 1.15 (for n = 949 individual deposition events). This level of dispersion is close to a Poisson-like process, which would have CV=1 and in which intervals are approximately exponentially distributed. Estimating the rate directly from the data with *λ̂* = 1/⟨*T*⟩λ=1/⟨T⟩ results in λ=0.09 and overlaying the corresponding exponential density *λ̂e*^−*λ̂T*^on the empirical interval distribution provides a good visual match. However, more stringent diagnostics indicate small but systemic deviations from an ideal homogenous Poisson process. A bootstrap Kolmogorov–Smirnov test rejects the fitted exponential model (D = 0.078, p = 2×10⁻⁴), the exponential Q–Q plot shows a mild upward deviation in the upper tail consistent with an excess of long intervals, and successive intervals exhibit significant positive serial dependence (lag-1 Pearson r = 0.18, p = 3.7×10⁻⁸; 891 adjacent pairs) (Supplemental Figure 4). Together, these results indicate that trail deposition is a stochastic process, close to a Poisson process, but also with a tendency to form bursts, which might be related to one nanotube decaying into several droplets.

### CSP trails are formed by stretching and collapse of membrane extrusions during gliding

Trails have been thought to be consisting either of pure protein fraction (*47*) or of membrane containing associated proteins (*36*). While we interpret our data to confirm the latter hypothesis, we also utilize a parasite expressing GFP anchored to the GPI of CSP as a marker for the plasma membrane (*42*). Migrating GFP-GPI parasites also showed trail depositions similar to those of CSP-GFP expressing parasites (Figure 3A). To investigate the possible membranous nature of the trail, we next used fluorescence recovery after photobleaching (FRAP) of tubules that were still connected to the sporozoite (Figure 3B). To this end we utilized oocyst-derived sporozoites as their migratory capacity was limited which allowed for easier bleaching and analysis. Bleached tubules recovered their fluorescence within a few seconds suggesting a connection to the sporozoite allowing a free flow of the protein from the parasite. Similar diffusive behavior was also seen within the trail deposited by a parasite expressing a variant of CSP-GFP where the GFP was introduced as in the GFP-GPI parasite (*42*) (Figure 3C).

**Fig. 3:**
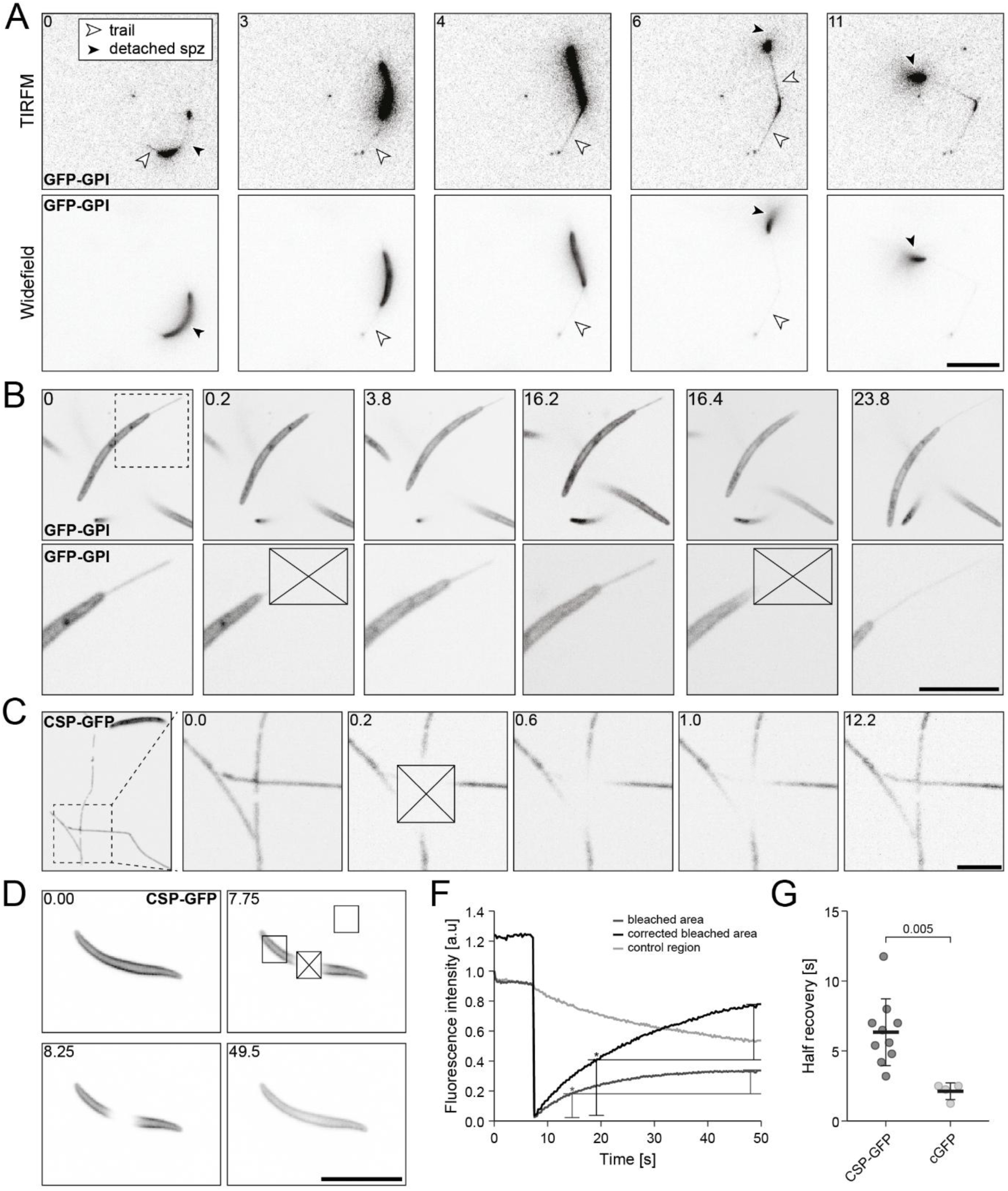
Investigating the membranous characteristic of sporozoites trails through TIRF and FRAP. **(A)** Regular TIRFM and the corresponding widefield fluorescence signal of a gliding salivary gland sporozoite expressing GFP with the GPI anchoring sequence of CSP (*42*). The sporozoite still shows formation of elastic tubules connecting the parasite to the surface (white arrowhead). The occasionally detached part of the sporozoite during stoppage is marked (black arrowhead). All scale bars 10 µm. **(B)** Repeated photobleaching of the extended trail in non-productive moving midgut sporozoite expressing GFP with a GPI anchor and subsequent recovery. The position of the zoom in (dashed square) and the photobleaching spot (crossed box) are marked. **(C)** A midgut sporozoite expressing a GFP-tagged version of CSP with the GFP positioned as in the GFP with GPI-anchor construct formed seemingly connected trails that recover fluorescence after bleaching. The position of the zoom in (dashed square) and the photobleaching spot (crossed box) are marked. **(D)** FRAP of a day 20 post infection midgut sporozoite expressing GFP-tagged CSP with the GFP between the repeat and C-terminal TSR domain resulting in defective egress (*42*). The FRAP spot (crossed box), the control region (left box) and background control region (top box) are marked. **(E)** Analysis of sporozoite in D. The mean fluorescence values of the areas are calculated for each time point in the series and normalized to t = 0. The corrected bleached area values are calculated by normalization with the control region to correct for general photobleaching through imaging. The time point of half recovery (*) is marked. **(F)** Half recovery times of sporozoites as in D and E compared to sporozoites expressing cytosolic GFP (*78*). Each point represents a single sporozoite and the standard deviation is shown. Significance is calculated through pair-wise comparison with Wild Type in Student’s t-test.

This high diffusability of CSP-GFP prompted us to investigate CSP-GFP on parasites. Again, we used oocyst-derived parasites for simpler analysis. Bleaching an area in the center of the sporozoite showed rapid recovery without noticeable directionality (Figure 3D, E). This suggests free diffusability of the CSP-GFP on the surface. Quantitative comparison with cytosolic GFP showed a half-recovery time t of the bleached CSP-GFP on the surface of around 6 to 7 seconds, while cytosolic GFP recovered in around 2 seconds (Figure 3F). This translates into estimates for the diffusion constant D=L^2^/p^2^t of 0.06 and 0.2 mm^2^/s, respectively, where L=2 mm is the size of the bleach spot (*55, 56*). This difference corresponds to the known diffusion difference between cytosolic and membrane environments and indicates that CSP interacts well with its membranous environment (*57*).

Together, these data suggest that trails are membranous deposits containing CSP. Trail formation is caused by a predominantly stochastic mechanism, instead of occurring in regular intervals, with consistent average frequency that is independent of gliding speed, yielding a strong correlation between deposition distance and speed, because the same material is distributed over a larger distance. The stochastic process is slightly bursty, which might correspond to single nanotubes being pulled out and decaying. As TRAP was previously found in the deposited trails (*32*) and the adhesion site turn-over as observed with RICM fits the characteristics seen with trail formation (Supplemental Figure 5), the turnover of adhesion sites may be an important element of how these nanotubes are formed and scissored.

### Imaging GFP-TRAP in gliding sporozoites reveals distinct adhesion sites and partial TRAP cleavage

TRAP is secreted from micronemes to the plasma membrane at the apical pole of sporozoites and thought to form distinct adhesion sites in conjunction with actin filaments that are in turn detached from the surface through cleavage with a rhomboid protease, presumably ROM4 (*32, 33*) (Figure 4A). Utilizing an endogenously GFP-tagged TRAP mutant line (*41*), we proceeded to perform live oTIRF imaging of gliding sporozoites expressing GFP-tagged TRAP. This showed distinct patterns of fluorescent signals, usually with a strong signal at the front of the sporozoite probably deriving from the micronemes, as well as an occasional smaller signal at the posterior, potentially deriving from the ER reaching into the TIRF field, both signals appeared fixed in position in respect to the moving sporozoite (Figure 4B). Dynamic imaging showed strong, consistent signals as the sporozoite circles, but further revealed weaker signals emanating from the apical side, moving along the sporozoite as the parasite migrated while staying fixed in respect to the substrate (Figure 4C, movies 6,7). As the parasites moved on, these patterns usually displayed the same profile of signal intensity (Supplemental Figure 6A). After an initial intensity peak, likely deriving from the strong apical signal, the intensity remains stable as the sporozoites moves over the site. The signal intensity only drops off as the site is left behind as part of the trail, after which the signal remains stable once again (Figure 4D, Supplemental Figure 6B).

**Fig. 4:**
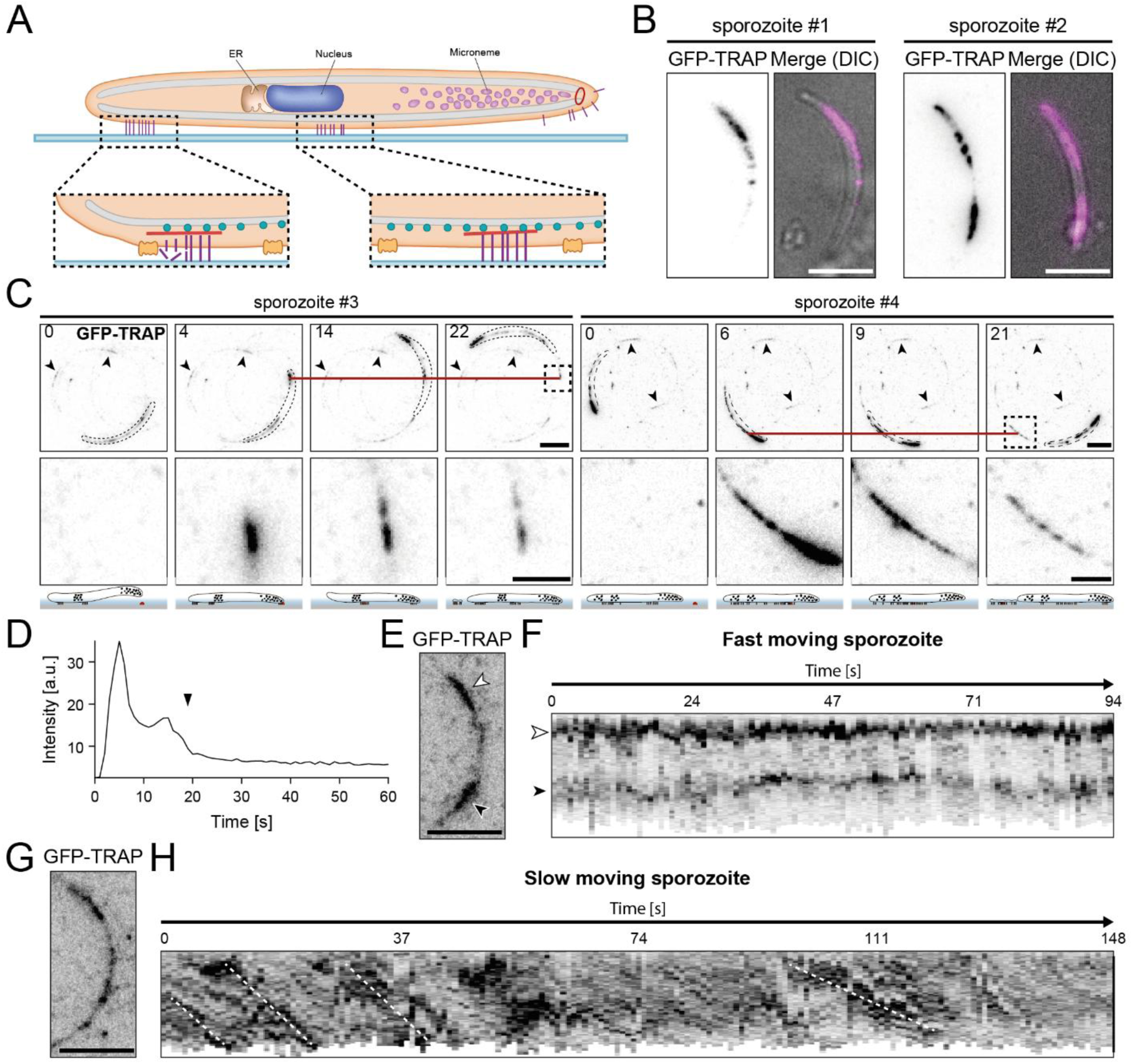
Visualization of TRAP adhesion sites and deposition as trails using oTIRF. **(A)** Expected presence of TRAP in the sporozoite. TRAP (purple) is stored in the micronemes, secreted at the apical tip, moved rearwards through actin-myosin and suspected to be cleaved at the posterior by a rhomboid protease (orange). **(B)** Exemplary localizations of GFP-TRAP in oTIRF and merged with DIC images. The apical tip is positions towards the top. Scale bar: 5 µm. **(C)** Turn-over of GFP-TRAP containing adhesion sites in gliding sporozoites visualized using oTIRF. The static position of exemplary adhesion sites over the imaging period (red line) and the position of the zoom in (dashed square) are marked. A schematic beneath the microscopy images illustrates the position of the sporozoite, adhesion sites and internal GFP-TRAP signal. The adhesion site in focus (red dot) and evanescent wave (blue) are marked. Scale bar: 5 µm and 2 µm for zoom in. **(D)** Summed fluorescence intensity of the marked adhesion site in C over the imaging period normalized to the intensity of the first frame. The moment the adhesion site is left as trail is marked (black arrowhead). **(E)** GFP-TRAP signal of a fast-moving sporozoite with the apical (white arrowhead) and posterior (black arrowhead) signal marked. Scale bar: 5 µm. **(F)** Kymograph of the GFP-TRAP intensity along the sporozoite in E throughout the imaging period. **(G)** GFP-TRAP signal of a slow-moving sporozoite. Scale bar: 5 µm. **(H)** Kymograph of the GFP-TRAP intensity along the sporozoite in G throughout the imaging period. The rough trajectory of exemplary, individual adhesion sites turned over through the sporozoites (white dashed lines) are marked.

Constructing kymographs from the oTIRF data revealed that the rapid movement is not visible in fast moving sporozoites, possibly because of the combination of low temporal resolution and weak adhesion sites (Figure 4E,F, movie 8). In fast moving sporozoites, only the stable signals of the GFP-TRAP from the presumed intracellular organelles were visible. In slow-moving sporozoites, however, the movement of the dynamic signals could be visualized, showing a similar turn-over pattern as previously described for presumed adhesion sites in RICM (*13*)) (Figure 4G,H, movie 9). Together, these data suggest that the trails visualized with the GPI-anchored CSP-GFP are surface-bound membrane fragments containing TRAP-based adhesion sites. These sites are seemingly not fully disassembled as the parasite moves on, but partially cleaved before being deposited as trails, similar to the situation in crawling cells (Figure 4I).

### Inhibition of TRAP cleavage results in reduced motility and altered trail deposition

In crawling cells, rear detachment occurs through a combination of biochemical cleavage of adhesion molecules and biophysical nanotube formation and rupture, which both have the effect of removing the constraint exerted by mature adhesions at the back. Together, these two processes allow the cells to tightly control rear detachment. To investigate the potential impact of rhomboid cleavage on the turnover of the adhesion sites in *Plasmodium* sporozoites, we next generated a parasite line that expressed a GFP-tagged version of a mutant TRAP, termed TRAP VAL, that cannot be cleaved efficiently by rhomboid proteases (*32*). The GFP was placed between the signal peptide and the first adhesive domain as was previously successful with wild type TRAP (Supplemental Figure 5A). Like the original TRAP-VAL line (*32*), GFP-TRAP-VAL parasites struggled to get into the salivary glands (Supplementary Figure 7), but those who did could be imaged. oTIRF microscopy readily revealed a large amount of surface TRAP in the GFP-TRAP-VAL mutant in comparison to the GFP-TRAP sporozoites (Figure 5A). GFP-TRAP VAL sporozoites migrated much slower, also in accordance with the TRAP VAL results (*32*), and the few motile parasites deposited large amounts of TRAP in the trails they left behind (Figure 5B, movie 10). To investigate the impact of rhomboid cleavage during motility, a more moderate inhibition was required. Therefore, we tested an inhibitor of rhomboid proteases, TLCK (*32*), in increasing concentrations and scored their motile behavior. At concentrations of 100 µM TLCK we found that sporozoite gliding was severely reduced, but a substantial fraction was still moving so that they could be further investigated (Figure 5C,D). Addition of 100 µM TLCK also showed seemingly stronger deposition of trails containing CSP-GFP (Figure 5E, movies 11,12), further supporting the link between the membranous deposits and non-cleaved adhesion sites. Tracking the dynamics of individual adhesion sites of wt GFP-TRAP showed a similar pattern of intensity and deposition (Figure 5F,G, movie 13). Interestingly, the loss of intensity from the stable plateau, as the parasite moved over the site, to the stable trail, remained the same between GFP-TRAP expressing sporozoites with and without TLCK treatment (Figure 5H). This suggests that the amount of TRAP left behind in the trail is not changed through the absence of proficient cleavage, and that the biophysical process of forming and collapsing nanotubes is sufficient to ensure rear detachment. However, the size and strength of the adhesion sites at the rear should strongly modulate this process. Rather than depositing larger adhesion sites after insufficient cleavage, the sporozoite may need to slow down or come to a halt before nanotubes can be pulled out of its rear.

**Fig. 5:**
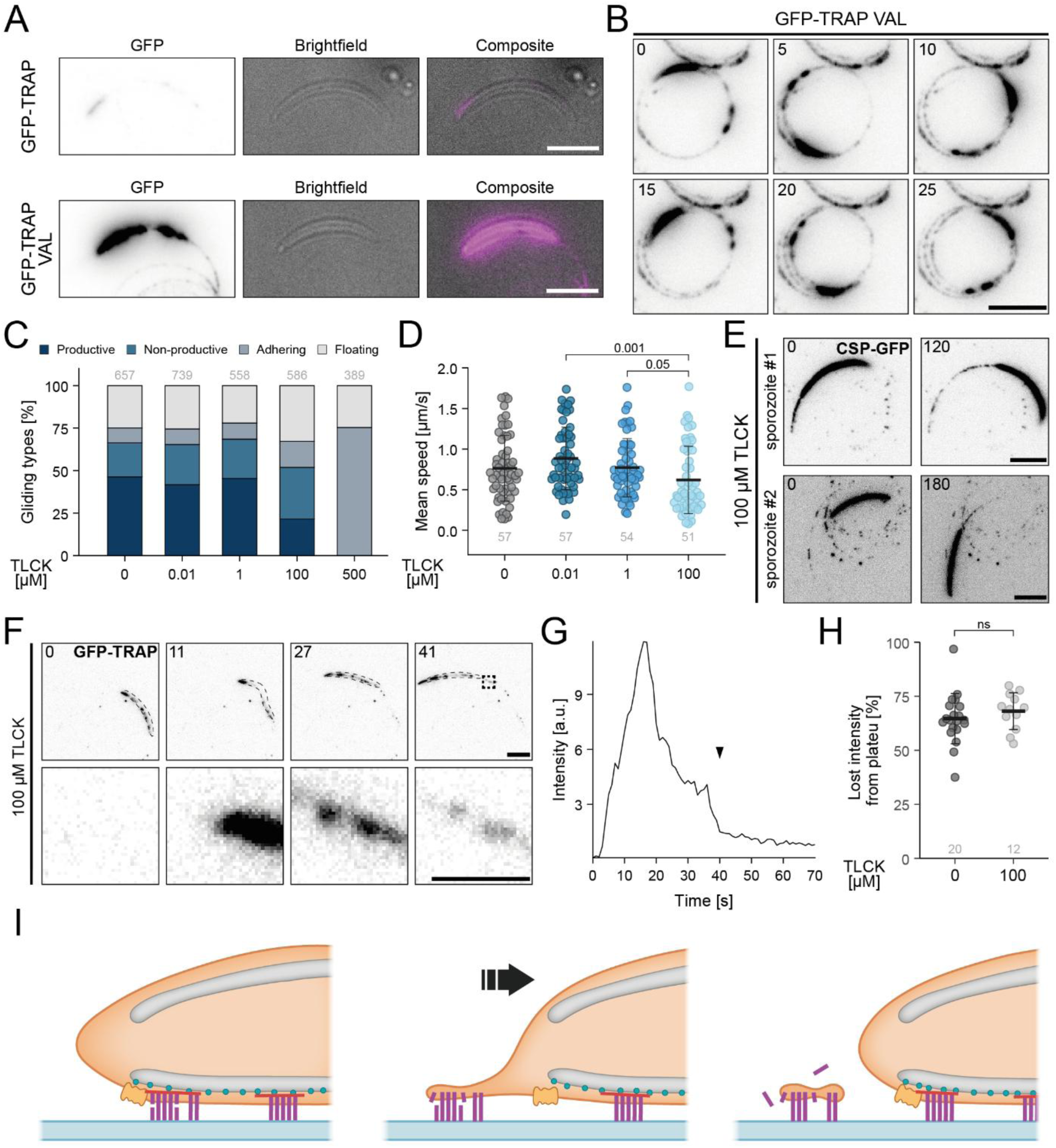
Inhibition of TRAP cleavage changes gliding and trail deposition behavior but does not result in an increase of trail signal. **(A)** oTIRF signal of a GFP-TRAP and GFP-TRAP VAL expressing sporozoite respectively. Parasites are shown in oTIRF, brightfield and a merge. Scale bar: 5 µm. **(B)** Time lapse of a gliding sporozoite expressing GFP-TRAP VAL in oTIRF. Scale bar: 5 µm. **(C)** Concentration dependent impact of TLCK treatment on gliding capabilities of sporozoites. Data consists of two replicates. **(D)** Mean speed of productively gliding sporozoites treated with different concentrations of TLCK. No productive moving sporozoites were present with a concentration of 500 µM. Significant differences in the whole population were determined via oneway ANOVA analysis. Afterwards, pair-wise comparison between all conditions with Student’s t-test is performed. The p-value is shown directly unless higher than 0.05. **(E)** oTIRF imaging of CSP-GFP expressing sporozoite treated with 100 µM TLCK. Shown are the first and last frame of the imaging period. Scale bar: 5 µm. **(F)** oTIRF time lapse of sporozoite treated with 100 µM TLCK gliding. The zoom in on an individual adhesion site is marked (dashed square). Scale bar: 5 µm. **(G)** Summed fluorescence intensity of the marked adhesion site in F over the imaging period normalized to the intensity of the first frame. The moment the adhesion site is left as trail is marked (black arrowhead). **(H)** Fluorescence intensity lost from the signal of the stable plateau before deposition to the trail after deposition in percentage with and without 100 µM TLCK treatment. Pair-wise comparison between the two conditions with Student’s t-test is performed. **(I)** Schematic of the proposed model of trail deposition. The TRAP-based adhesion sites reaching the posterior end are partially dissolved before being ripped out with some membrane deposits as the parasite moves on.

To further investigate the relation between adhesion site disassembly, deposition and parasite speed, we plotted the fluorescent intensity of individual adhesion sites in GFP-TRAP parasites with and without TLCK treatment as well as the instantaneous speed. We opted to plot the speed as averages over a 5 second interval to smooth speed variations (Figure 6). This showed that TLCK treated sporozoites still reach similar speed peaks as non-treated parasites. The recovery of the speed after temporal slowing from trail deposition, however, appeared different. In the moment after the first adhesion site deposition as trails in control sporozoites, the speed recovered to almost its peak over the next seconds. Only when multiple adhesion sites overlapped, the speed recovery is slower (Figure 6A, movie 14). After treatment with TLCK, the speed did not recover to its full peak and additional following adhesion sites result in overall slowing down of the parasite (Figure 6B, movie 15).

**Fig. 6:**
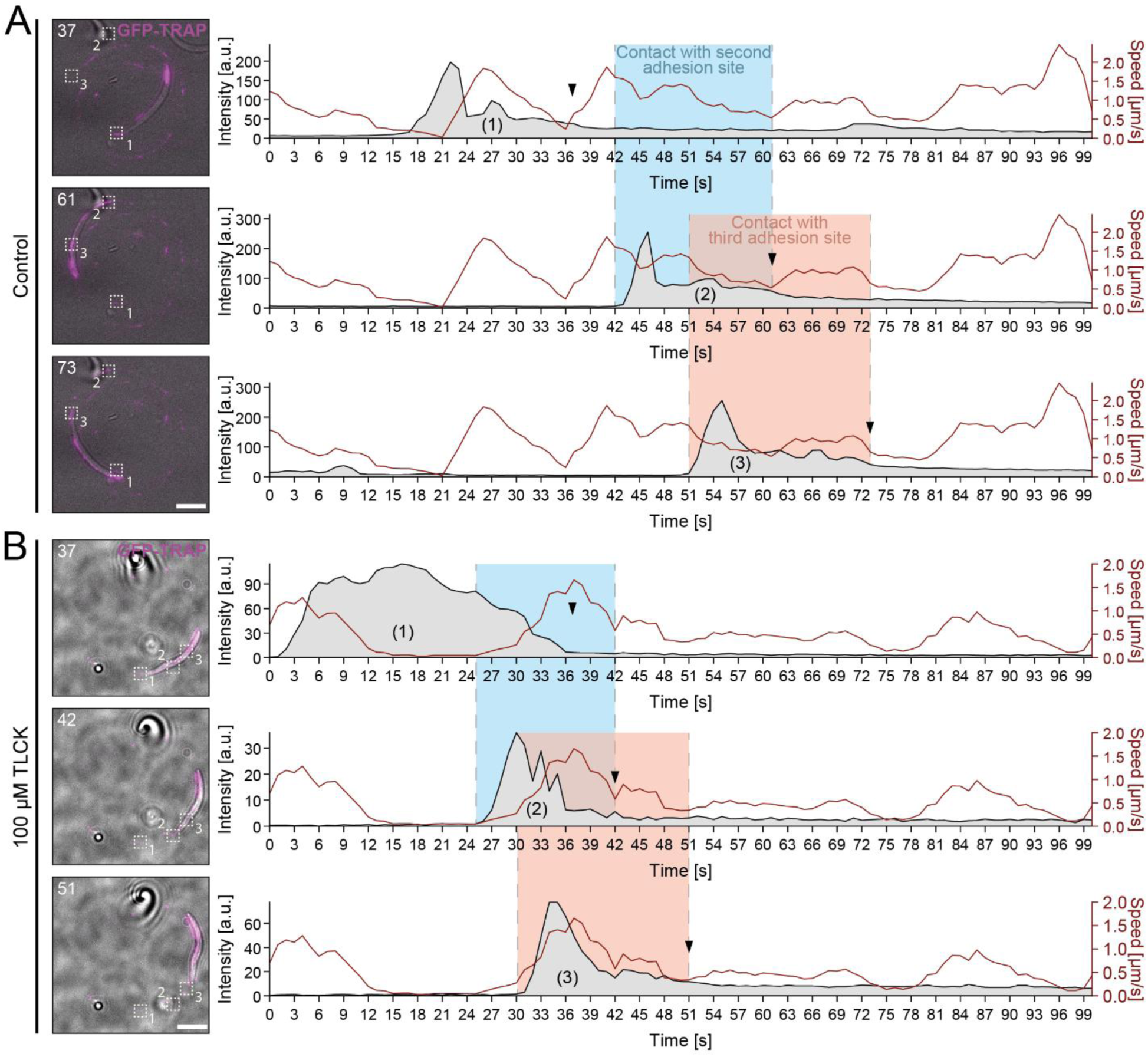
Increased trail deposits after TLCK treatment slow sporozoites down. Adhesion analysis of an **(A)** untreated and **(B)** 100 µM TLCK treated sporozoite. The summed fluorescence intensity of the marked adhesion sites (1, 2, 3) are shown alongside the instantaneous speed of the sporozoite over the imaging time. The instantaneous speed is calculated as a mean over a 5 s window. The moment of deposition as trail is marked (black arrowhead). The position of each chosen adhesion site is marked (dashed square) in a GFP-TRAP oTIRF and brightfield merge. The time frame in which the sporozoite is engaged with the second (blue area) and third (red area) of the three analyzed adhesion sites are marked. Scale bar: 5 µm.

### Distinct actin networks in sporozoites treated with TLCK

To investigate the underlying motor machinery of the glideosome, we sought to image the dynamics of actin filaments. Expressing an actin chromobody in sporozoites revealed distinct localization of actin filaments with a majority of sporozoites showing filament accumulations at the rear end (*46*). Imaging these sporozoites with the selective excitation of oTIRF microscopy revealed a higher resolution image of these filaments and unraveled a helical structure that stretched several micrometers from the rear end of the sporozoite towards the center of the cell (Figure 7A, movies 16,17). After treatment with 100 µM TLCK, sporozoites displayed a small, dot-like signal instead and only a minority still featured a longer filamentous actin localization (Figure 7B,C, movie 18). The dot-like accumulation was concentrated at the very rear with a radius of less than 500 nm (Figure 7D). On rare occasions we also found actin filaments (i.e. chromobody) deposited in the trail (Figure 7E, movie 19) further suggesting that trails are membrane deposits that are surface-attached through intact adhesion sites and containing cytoplasm.

**Fig. 7:**
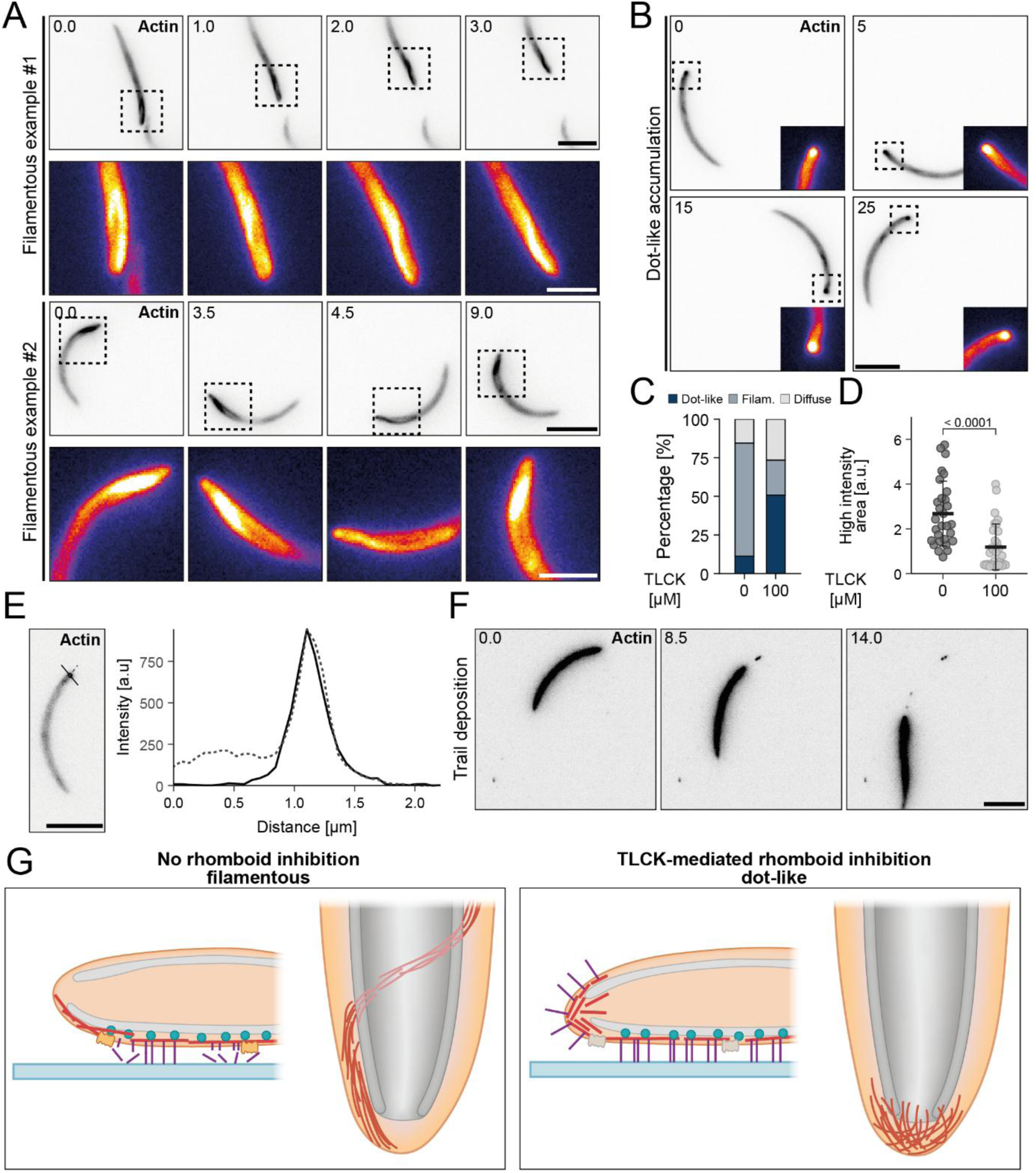
Inhibition of TRAP cleavage disrupts actin structures in gliding sporozoites. **(A)** Examples of filamentous actin accumulations at the posterior of sporozoites expressing actin chromobody (ChrB)-GFP gliding in oTIRF. The position of the zoom in (dashed square) is marked. Scale bar: 5 µm and 2 µm for zoom in. **(B)** Example of dot-like actin localization in an actin ChrB-GFP expressing sporozoite treated with 100 µM TLCK. Position of the zoom in colored with a different lookup table is marked (dashed square). Scale bar: 5 µm. **(C)** Prescence of the different actin localization patterns determined through blinded classification in gliding sporozoites with and without 100 µM TLCK treatment. **(D)** Area of pixels with actin ChrB-GFP intensities higher than 80% of the maximum intensity value for sporozoites with and without 100 µM TLCK treatment. Pair-wise comparison between the two conditions with Student’s t-test is performed. **(E)** Fluorescence intensity profile of the dot-like actin accumulation along (dashed line) and perpendicular (solid line) of the sporozoite axis. Scale bar: 5 µm. **(F)** Example of actin ChrB-GFP containing trails deposited during gliding motility in a 100 µM TLCK treated sporozoite. **(G)** Schematic illustration describing the possible explanation for the observed accumulation patterns. Without inhibition of TRAP cleavage, actin can disassociate from bound TRAP and accumulate along the parasite length in the larger filamentous bundle. Defective rhomboid cleavage of TRAP may result in actin remaining bound to TRAP and accumulate at the most posterior end. Potentially inhibiting further actin depolymerization.

Taken together these data show actin filaments to accumulate in elaborate, helical bundles during motility and that the cleavage of TRAP and the disassembly of adhesion sites also impacts the underlying actin filaments. The helix-like accumulation may be caused by large numbers of actin filaments, disassociated from the adhesion sites and not anymore involved in active motility, piling up to form a longer actin bundle that is forced into a helical form through the spatial restrictions of the underlying IMC (Figure 7F). If TRAP cleavage is inhibited, the actin filaments may be pulled further to the posterior end, unable to disassemble as the uncleaved TRAP remains attached, forming the more condensed dot-like accumulation.

## Discussion

Investigation of the formation and dynamics of adhesion sites in the rapidly migrating apicomplexan parasites have so far been hampered by the lack of appropriate surface sensitive imaging methodology. Prior work has examined adhesion dynamics by reflection interference contrast microscopy, RICM (*13, 15, 24*). In these studies, the presence of adhesion sites was inferred from the close apposition of the parasite to the glass surface as revealed in the interference pattern. Only one study used TIRF microscopy to investigate *Plasmodium* sporozoite adhesion to the substrate (*58*). This revealed a stepwise attachment of the sporozoite to the substrate prior to gliding. In that study, we however failed to provide insights into adhesion dynamics during gliding due to the inhomogeneous evanescent field resulting in orientation-dependent uneven fluorescence signals. The use of oTIRF allowed the generation of a homogeneous evanescent field and hence the reliable imaging of adhesion sites in gliding sporozoites. This allowed us to image the formation of TRAP containing concentrations on the parasite surface that are deposited on the substrate as the parasites move forward, hence suggesting that these are adhesion sites used by the parasite to move forward. Interestingly, approximately 40% of the fluorescent GFP-TRAP signal (of the “adhesion site”) is deposited on the trail. This could indicate that only about half the TRAP in an adhesion site is cleaved by a rhomboid protease and disperses in the medium. Alternatively, more of the TRAP may be cleaved but could remain with the adhesion site by binding the surface via the A-domain. Mutational inhibition of rhomboid cleavage led to an accumulation of GFP-TRAP VAL on the surface, as previously described, and to stronger trails (*32*). However, the strong effect on continuous motility did not allow for detailed investigation of adhesion site dynamics.

Concentration-mediated chemical inhibition of rhomboid (and possibly other) proteases through the serine-protease inhibitor TLCK showed similar but less severe effects on motility as mutational inhibition. Curiously, this inhibition still resulted in the same relative amount of TRAP being deposited in trails as in untreated sporozoites, indicating that only once a certain amount of TRAP is removed from an adhesion site, it can be deposited as trail and the parasite can move on. Insufficient cleavage of TRAP therefore, may result in the parasite being in contact with posterior adhesion sites for longer and therefore slowing down fitting to observations with traction force microscopy, where a stalling force is found at the rear in sporozoites that are temporarily stuck at the rear end (*13, 59*). Alternatively, or in addition, inhibition of adhesion site disassembly may result in overall more adhesion sites persisting. In untreated sporozoites these adhesion sites may be disassembled before reaching the posterior end.

TRAP is thought to link to actin filaments and being moved rearwards by myosin motors generating a retrograde membrane flow of adhesins (*60*). This retrograde flow is faster than sporozoite forward movement, suggesting that adhesion site formation slows actin retrograde flow. Curiously, the retrograde flow is faster in the absence of the TRAP like protein TLP (*60*), suggesting that adhesion sites are modulated by their composition. Retrograde flow is also faster if actin filaments are stabilized by small concentrations of jasplakinolide (*60*), suggesting a complex interplay between different types of adhesins and actin filaments for adhesion site formation, force generation and adhesion site turnover. Actin filaments were only recently visualized by cryo-electron tomography in sporozoites (*61*) and other parasites (*62*), but not associated with adhesion sites or during motility. How adhesion sites are structured is thus still an important open question to answer. oTRIF microscopy coupled with fluorescently labelled surface and glideosome proteins will likely play a major role in dissecting adhesion site composition, formation and turnover. The ability of directly visualizing the dynamics of adhesion sites through oTIRF opens the doors to investigate the process of gliding motility in other *Plasmodium* stages such as ookinetes or merozoites as well as other apicomplexan parasites such as the tachyzoites of *T. gondii*.

Interestingly, we observed different forms of actin filament accumulations at the rear end of sporozoites: dot-like and helix-like assemblies using oTRIF microscopy of parasites expressing an actin filament recognizing chromobody. Previous widefield fluorescence microscopy could not distinguish these different assemblies (*46*). While the helical assemblies dominated in the wild type, parasites treated with the rhomboid inhibitor TLCK showed mostly the dot-like assemblies. This suggests that the arrangement and/or turnover of actin filaments within the sporozoites is modulated by the state of the TRAP proteins spanning the plasma membrane. Natural cleavage of TRAP proteins, as in wild type parasites, led to the helical accumulations of actin filaments at the rear associated with optimal motility, while non-cleaved TRAP, associated with slow motility, led to the much smaller dot-like accumulations (Figure 6F). Intriguingly, this concentrated accumulation is similar to the accumulation of F-actin in *T. gondii* tachyzoites which may indicate differences in adhesion capabilities of adhesins in the two parasites (*63*). This seems at first to be counter-intuitive as we would have expected a faster actin filament turnover and hence smaller accumulations in the non-TLCK treated parasites. Yet, it cannot be excluded that the arrangement of actin filaments at the rear might play a part in pulling the very rear off the substrate, as is the case in crawling cells (*64*). We speculate that this helical arrangement could be achieved by the detachment of actin filaments from TRAP enhanced by proteolytic cleavage of TRAP. The now ‘free floating’ actin filaments might arrange differently to accommodate newly arriving filaments from the front, while non-detached actin filaments might simply accumulate in the back and hamper migration. Cryo-electron tomography of the two conditions might reveal interesting details of the different arrangements as will the expression of the chromobody in parasite lines featuring mutated adhesins or actin binding proteins.

Visualizing the deposits of surface proteins has been a well-established way to quantify gliding motility of apicomplexan parasites without live microscopy. Yet, its relevance for motility *in vivo* is largely unclear. While CSP was detected on infected salivary glands (*65*), it was not yet visualized in the path of a migrating sporozoite *in vivo*. Using the *in vitro* polyacrylamide gels gliding assays we could detect CSP via immunofluorescence methods as regular dot-like signals in helical paths, similar to the paths of gliding sporozoites in the gels (*10, 51*) suggesting that trail formation could also occur *in vivo*. The mechanisms behind the formation and composition of these trails are, however, poorly understood. It was believed that these trails are formed through shedding of proteins onto the surface in the absence of membrane deposition (*47*). The fluorescent tagging of the GPI-anchored CSP allowed oTIRF imaging of the protein as a proxy for the membrane. The stretching and collapse events visualized suggest the trails to form through whole membrane tubules that fragment into smaller vesicles at some substrate-bound anchoring points. Such a process is well-known as the Rayleigh-Plateau instability for fluid cylinders under tension, as known from water faucets, whose cylindrical water columns decay into droplets, because this lowers surface energy at the same volume. The Rayleigh-Plateau instability has been implicated in many biological processes, including the pearling of cellular protrusions under weakening of the cytoskeleton (*54*) and pearling of neurons after changes in osmotic pressure (*66*). Very recently, this biophysical mechanism has also been implicated for shape control of mitochondria (*67, 68*). Because the Rayleigh-Plateau instability is mainly driven by surface tension, it would be highly interesting to measure its values in moving sporozoites, as done earlier for moving keratocytes. For keratocytes, it has been shown that surface tension increases with increased actin polymerization and increased adhesion strength (*69, 70*). Interestingly, keratocytes might share with sporozoites the features that they have few membrane reservoirs, different from e.g. fibroblasts. This suggests that surface tension might be higher at the sporozoite rear, where actin accumulates and mature adhesion might not be sufficiently weakened by cleavage. This then might increase the instability, thus making sure by physical means that the sporozoite can move forward by leaving behind its adhesion sites as droplets deposited in the trail. We note that sporozoites are strongly primed to reach blood vessels, and that a new stage starts in the liver, so it does not matter if material is lost in the skin.

Our deposition model for *Plasmodium* also fits with lipids and cytoplasmic proteins that were shown to be present in some trails of *T. gondii* (REFs) as well as the occasional dots of actin we could visualize in some trails. The vesicle formation itself could be an active process or result from wound healing after physical disruption, potentially explaining previous findings of cytoplasmic proteins on the sporozoite surface (Refs: Lindner surface proteome). The intriguing relationship between the rate and distance of trail deposits suggests a mechanism that is on one hand tightly coupled to the speed and therefore the gliding machinery of sporozoites, while also having a random or even burst-like periodicity independent of the speed. The time-independent distribution of trail events combined with the presence of TRAP-containing trails fit a model of non-cleaved adhesion sites as foundation of trail deposits.

In order to truly understand apicomplexan gliding motility and adhesion dynamics the obvious next step is to analyze other motile *Plasmodium* stages, mainly ookinetes, and *T. gondii* tachyzoites with oTIRF to investigating the role of their respective mayor surface proteins (P25&P28 and Sag1) as well as their main micronemal adhesin (CTRP and MIC2) (*71–74*). With dual color oTIRF, both the mayor surface molecule as well as the adhesin could be imaged simultaneously, especially in the much slower ookinetes but potentially also in sporozoites with altered motility as observed in the *hsp20*(-) sporozoites (*75*). So far, trail formation has evolved from the apparent artifact that allows for a static gliding assay proxy to a deeper understanding how sporozoites establish, turn over and disassemble their adhesion sites during the rapid gliding motility.

In summary, our data show the turn-over of adhesion sites and their deposition as trails likely through the elongation and collapse of membrane tubes that are anchored to the substrate by TRAP. This contrasts the long-held view that trails consist mostly of surface proteins that are shed through some active process (*47*). However, many aspects of the interplay between adhesion site turn-over, TRAP cleavage by rhomboid protease and CSP function in gliding remain open. Answering those will increase our understanding of a fundamental biological process and might yield new insights into how the sporozoite can be stopped.

## Materials and Methods

### Animal work

All mice experiments were performed according to the FELASA and GV-SOLAS standard guidelines and were approved by the German authorities (Regierungspräsidium Karlsruhe). All parasite lines were generated in the *P. berghei* ANKA parental strain and maintained through infection of 4 -6 week old female CD1 mice. Transfections and clonal lines were performed and selected for as previously described (*43*). Mosquitos were infected by initial injection of a naïve mouse with frozen parasite stock and after a sufficient parasitemia was reached, the mouse was anesthetized and the blood was harvested through heart puncture. 20 million parasites were transferred into two naïve mice and after 3 to 4 days, exflagellation of gametes was confirmed via microscopy. The mice were anesthetized and female *Anopheles stephensi* (strain SDA 500) were allowed to feed for 20 to 30 min. The mosquitos were then kept at 21 °C.

### Generation of CSP-GFP expressing parasite line

The plasmid to generate parasite lines expressing CSP-GFP as an additional copy was previously generated (*42*). Alternative digest of the plasmid with EcoRI and EcoRV allowed double homologous recombination of the CSP-GFP construct into chromosome 12. Correct integration and clonality was confirmed through the presence of the correctly sized 5’ (P134 and P244) and 3’ (P135 and P137) and absence of the WL wild type (P134 and P137) bands in genotyping PCR amplification.

### Generation of GFP-TRAP VAL expressing parasite line

The plasmid with the cleavage site mutation VAL (AIGGIIGG => VALGGIIGV) was a kind gift of Photini Sinnis (*32*). We previously inserted a GFP between the signal peptide and the A-domain (after the first 35 amino acids, flanked by a 6 amino acid linker upstream and a 12 amino acid linker after the GFP (*41*). Prior to transfection the plasmid was linearized with EcoRI and KpnI and integrated into the TRAP locus, replacing the wt TRAP.

### Determining oocyst numbers in infected mosquito midguts

Between 10 and 12 days post infection, the midguts of infected mosquitos were collected and incubated in 1% Nonidet P-40 in PBS for 20 min at room temperature. Afterwards, midguts were stained in 0.1% mercurochrome in PBS for 1 h at room temperature and washed thrice with PBS. The mdiguts were transferred to a glass slide, covered with a cover slip and sealed with paraffin. Oocysts were imaged and counted using an inverted Axiovert 200 M microscope from Zeiss, GFP-channel and a 10x objective.

### Determining the number of sporozoites in mosquito organs

Between day 18 and 21 post infection, the salivary glands and midgut of infected mosquitos were collected in PBS. After smashing the organs, the solutions were diluted and the number of sporozoites were counted using a hemocytometer and a widefield microscope using a 40x phase contrast microscope. The number of sporozoites per infected mosquito were collected for the salivary glands and midguts respectively.

### Sporozoite motility assay

Sporozoites were isolated from infected mosquito salivary glands after at least 18 days post infection. The glands were crushed and diluted in BSA in RPMI for a final concentration of 3% BSA. The sporozoite solution was transferred into an optical 96-well plate and centrifuged at 800 g for 3 min. When determining the influence of inhibitor addition on gliding motility, a serial dilution of TLCK in 3% BSA in RPMI was prepared. TLCK is added in 1:2 dilutions to the sporozoite solution after centrifugation for final concentrations of 0 µM, 0.01 µM, 1 µM, 100 µM and 500 µM.

All sporozoites were imaged using a widefield microscope in the DIC channel for 100 time points and a frame rate of 3 s. Motility was scored according to classification into productively motile, non-productively motile, adhering and floating sporozoites. All parasites that moved at least a full circle and were continuously moving for half the imaging period were scored as productive. All other sporozoites displaying some form of motility were scored as non-productive. Sporozoites that remained continuously above the surface in media were defined as floating. Parasites that did contact with the surface but did not move were categorized as adhering. Gliding speed was determined by tracking individual sporozoites for at least 50 time points using the manual tracking plugin of Fiji. All motility assays were blinded prior to analysis. Imaging was performed at an inverted Axiovert 200 M microscope from Carl Zeiss Microscopy with a 25x water-immersion objective (0.8 NA), PRIME BSI R-M-16-C camera and a frame rate of once every 3 s for 100 time points in DIC.

### TIRF gliding assay

Sporozoites were isolated from infected mosquito salivary gland after at least 18 days post infection. For this, the glands were crushed and carefully underlaid with 17% Accudenz in ddH_2_O (*76*). The gradient is centrifuged at 2500 RPM for 20 min without breaking. The sporozoites are collected at the interphase between media and Accudenz, centrifuged again and resuspended in 3% BSA in RPMI after removal of the supernatant.

For oTIRF imaging, the sporozoites were transferred into an 8-well glass-bottom dish that was previously rinsed with PBS. For oTIRF imaging under TLCK conditions, TLCK in 3% BSA in RPMI was added for a final concentration of 100 µM. The dish was centrifuged at 800 g for 3 min and gliding sporozoites were imaged in the DIC and GFP channel. Time series were taken for 200 time points with an exposure time of 100 ms and a frame rate of 1 s for GFP-TRAP and anti-actin-ChrB expressing sporozoites and 0.5 s for CSP-GFP expressing sporozoites. TIRF imaging was performed on an inverted Zeiss Axio Observer 7 equipped with a Visitron Systems ORBITAL Ring-TIRF (oTIRF). The GFP labelled parasites were imaged with a 488nm (100mW) Laser under TIRF illumination angle using a Chroma Quadband 405/488/564/640 TIRF filter set. For transmitted light images, a CoolLED pE-100 was used. All images were acquired with a PRIME-BSE-Express back-illuminated sCMOS (6.5µm dexel size, final pixel size of the image 0.065µm/ pixel) using Visiview 5 microscope-control software (Visitron).

### FRAP analysis

Sporozoites were isolated from infected midguts after at least 18 days post infection. The tissue was crushed in 3% BSA in RPMI, transferred to an imaging dish and centrifuged prior to imaging.

For FRAP, movies were taken with 4 pfs and an area of 20 x 20 px or 25 x 25 pixel was bleached (PK cycles = 5, PK step size = 1, Spot period = 50, PK Spot Cycles = 15-20, PK spot size = small. Additionally, the lasers 405 nm, 440 nm, 488 nm, 514 nm, 561 nm and 640 nm were set to 100%.

For analysis, mean fluorescent signal was calculated for areas of identical size including the bleached area (b), a control area (c) of the same sporozoite and a background area (n). The corrected bleached value (f = (b - n)/(c - n)) was calculated for each frame alongside the minimum (min) and maximum (max) intensity value reached after bleaching. From this, the half recovery time t_1/2_ was calculated as the time point from bleaching until the half recovery value I_1/2_ =(max-min)/2 was reached. The second frame after t_1/2_ was exceeded, was used to calculate the time of half recovery. FRAP microscopy was performed using the PerkinElmer Nikon spinning disc confocal with the Volocity FRAP software using the UltraVIEW FRAP unit.

### Immunofluorescence staining after 3D motility in polyacrylamid hydrogels

Sporozoites were allowed to move in soft polyacrylamide hydrogels with 3% acrylamide and 0.03% bisacrylamide as described in (*10, 51*) for 1 h. Afterwards the gel was immersed in 4% PFA in PBS for 1 h at room temperature. The gel was washed briefly 3 times by removing the excess PFA solution and adding PBS. The final washing step was extended to 1 h. 3% BSA in PBS was added and the gel was blocked at 4°C overnight. Primary antibody solution (1:300 µg/µL 3D11 monoclonal a-CSP antibody in 3%BSA in PBS) was added and incubated for 8 h at room temperature. After washing thrice in PBS for 10 min, secondary antibody solution (1:400 AF594-a-mouse in 3% BSA in PBS) was added and incubated at 4 °C overnight. The gel was washed 3 times in PBS for 10 min and a second cover slip was added and sealed with paraffin. Fluorescence was imaged with the PerkinElmer Nikon spinning disc microscope and an EM-CCD Hamamatsu camera. Z-stacks were created with a slice size of 1 µm.

### Western Blot analysis of CSP in salivary gland sporozoites

At 17 days post infection, salivary glands of 30 PbANKA and CSP-GFP infected mosquitos were collected and smashed. Sporozoites were lysed in RIPA buffer with added protease inhibitor for 1 h on ice and stored at -80 °C until further use. Samples were diluted with 4x Laemmli sample buffer, cooked at 95 °C for 5 min, briefly centrifuged and loaded on a precast Mini-PROTEAN TGX 4-15% polyacrylamide gel (Bio-Rad). The sample was separated for approximately 45 min at 200 V and semi-dry blotted onto a nitrocellulose membrane. The membrane was blocked in 5% milk powder in TBS-T for 1 h at room temperature. Afterwards, the primary antibody (Roche mouse-αGFP (Ref#: 11814460001)) was added 1:1,000 in 5% milk powder in TBS-T and incubated over night at 4°C. The membrane was washed three times in TBS-T for 10 min. Afterwards, the secondary antibodies (-IRDye 800CW-αmouse) was added in a dilution of 1:10,000 in TBS-T and incubated for 2.5 h at room temperature. After washing trice with TBS-T, the fluorescence signal was measured at a LI-COR Odyssey CLx. The membrane was stripped for 20 min at 56 °C under harsh conditions (62.5 mM TRIS pH = 6.8; 2% SDS; 100 mM β-mercaptoethanol). Afterwards the membrane was blocked, incubated with the second primary antibody (3D11 mouse-αCSP (*18*)) and imaged as described above.

### Analyzing trail deposition events in oTIRF data through generation of kymographs

The distance and interval between trail deposition events was analyzed through a custom FIJI macro. The sporozoite fluorescent CSP-GFP signal in oTIRF was tracked manually by defining the posterior position of the sporozoite in each frame. A segmented line is generated along every third sporozoite position to remove smaller deviations emerging from very short distances moved and avoid artificially increased track distances. The fluorescence intensity at each position of the segmented line and their 4 orthogonally neighboring pixels is measured and the mean is calculated, resulting in a fluorescence intensity profile along the sporozoite path. This measurement is repeated for each frame of the time series and the profiles are stacked vertically, resulting in the final kymograph. The tracking points are also utilized to calculate the instantaneous and the mean speed of every analyzed sporozoite. The trail deposition events are identified in the kymograph as the original intersection of the vertical trail and the diagonal sporozoite signal. Their position is manually determined and the distance and interval to the previous trail event are calculated using the frame rate and pixel value of the time series.

### Quantification of actin accumulation

Differing actin accumulation in gliding sporozoites expressing the actin-ChrB was characterized by blinded classification into filamentous, diffuse or dot-like structures. Furthermore, quantitative analysis was performed using a custom Fiji macro that measures the area of pixels within the TIRF imaging of gliding sporozoites that achieved above 80% of the highest pixel intensity of the sporozoite. The mean measure of three frames of each sporozoite was calculated and displayed. Claude (architecture version Sonnet 4.6, Anthropic 2026) was used as coding assistance in RStudio.

### Image processing and illustration

Image analysis was performed with FIJI (LOCI, Wisconsin-Madison, USA) (*77*) and MatLab (MathWorks Inc., Massachusetts-Natick, USA). Figures were generated with Adobe Illustrator 2020 (Adobe, München, Germany). Graphs were generated with RStudio (Posit Software, Massachusetts-Boston, USA). Images and movies were individually adjusted for contrast. Fluorescence oTIRF images in figure 5A were adjusted to the overexposed GFP-TRAP VAL parasite.

## Acknowledgements

We thank Miriam Reinig and her students for help in mosquito rearing, Shane Scott for providing software for kymograph analysis, Photini Sinnis, Markus Ganter and Franziska Hentzschel for comments on the manuscript and helpful discussions, as well as Photini Sinnis for the TRAP-VAL construct. KW is a member of the Heidelberg Biosciences International Graduate School (HBIGS). We acknowledge the microscopy support from the Infectious Diseases Imaging Platform (IDIP) at the Center for Integrative Infectious Disease Research. The *Plasmodium* database PlasmoDB facilitated this work.

## Disclosure of artificial intelligence tools use

ChatGPT (architecture version GPT 5.2, OpenAI 2025) and Claude (architecture version Sonnet 4.6, Anthropic 2026) were used as coding assistance in RStudio and FIJI for generation of macros.

## Funding

This project was funded by grants from:

Deutsche Forschungsgemeinschaft (DFG, German Research Foundation) SFB 1129 “Integrative analysis of replication and spread of pathogens”, project number 240245660.

DFG SPP 2332 “Physics of Parasitism” (FR2140/13-1; and FR2140/13-2).

Wellcome Trust Discovery Award (225844/Z/22/Z).

Humboldt foundation postdoctoral fellowship to CTM.

DFG CellNetworks Cluster of Excellence.

The funders had no role in study design, data collection, and interpretation or the decision to submit the work for publication.

## Author contribution

Conceptualization: KW, MS, FF.

Methodology: KW, MS, FF.

Investigation: KW, MS, YSA, CTM, SO.

Formal analysis: KW, MS, LL, MCU.

Visualization: KW, MS, LL.

Supervision: CSU, USS, VL, FF.

Writing-original draft: KW, MS, FF.

