## Supplemental figures 1 to 7 for "Connecting adhesion dynamics and trail formation in malaria parasites by imaging the major surface antigens CSP and TRAP"

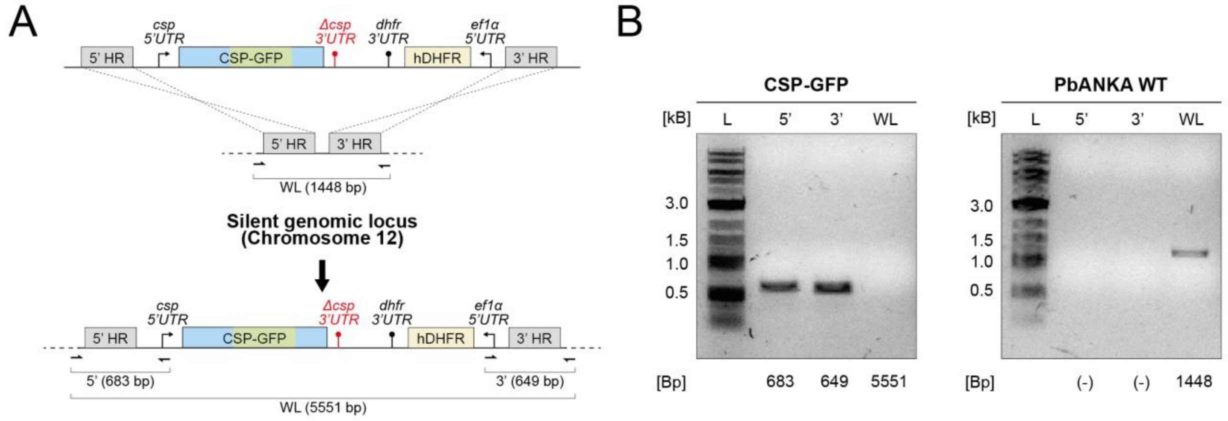


**Fig. S1.**
**(A)** Integration strategy for the generation of the CSP-GFP line consisting of a GFP integrated between the repeat and C-terminal region of CSP. The tagged version of the protein is integrated into a silent genomic locus on chromosome 12 via two homology regions flanking the CSP-GFP and the selection marker. The CSP-GFP is expressed under the 5' and 3' UTRs of csp with the 3'UTR being shortened to 286 bp. The selection marker consists of the human dihydrofolate reductase (hDHFR) under the ef1α 5' UTR and the *P. berghei* dhfr 3'UTR. Indicated are the primer binding sites and expected sizes of PCR amplicons for the genotyping in B. **(B)** Genotyping PCR of the final CSP-GFP clone utilized for this study and a parental PbANKA Wild Type control. The ladder (L), 5' band (5'), 3' band (3') and whole locus (WL) are indicated. The expected sizes are shown below.

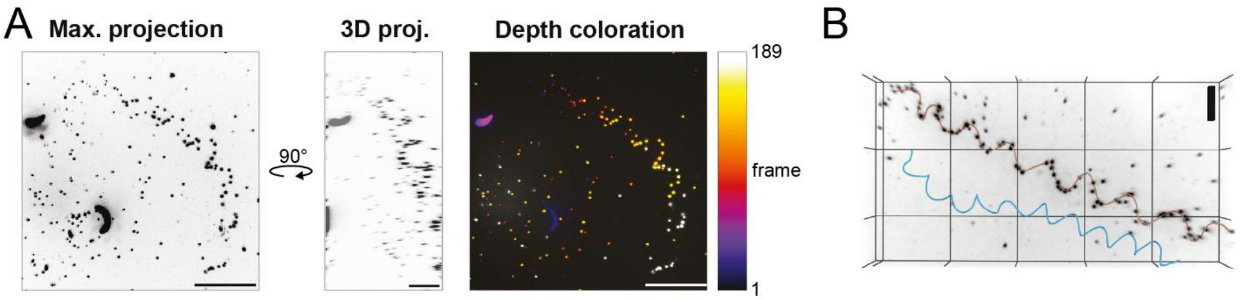


**Fig. S2.**

**(A)** A second example of Immunofluorescence of CSP in hydrogel environment after sporozoite motility using a spinning disc microscope. Shown is the maximum projection of the z-stack as well as a 3D projection, rotated for 90°. A depth coloration as an additional visualization of the 3D structure. Scale bars: 20 µm in maximum projection and depth coloration; 10 µm in 3D projection. **(B)** Fitting of a helical path along the CSP deposits in 1 H (red line). For comparison, the tracked path of a sporozoite moving in a 3D hydrogel of the same composition is shown (blue line) *(10)*.

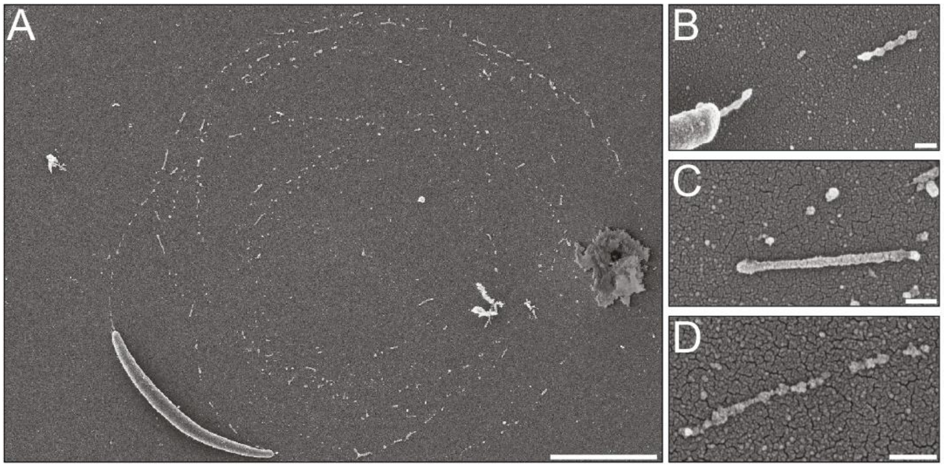


**Fig. S3.**

Scanning electron microscopy of a sporozoite and trails left after gliding motility. **(A)** An overview of a sporozoite and circular trails likely left behind by this specific sporozoite. **(B-D)** Trails from other sporozoites showing distinct stages of pearling. Scale bars: 200 nm.

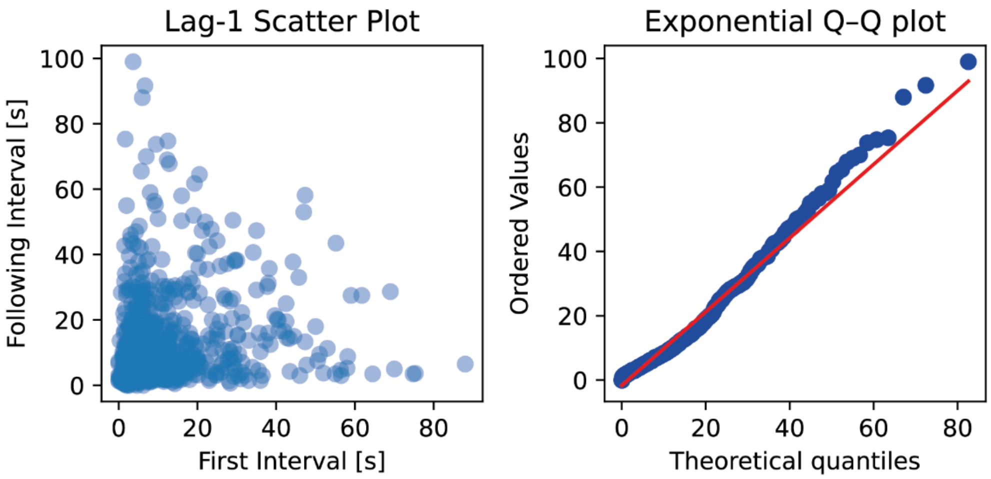


**Fig. S4.**
The intervals are plotted versus the interval following directly afterwards in a Lag-1 scatter plot. Q–Q plot of trail deposition intervals against a fitted exponential model. Observed inter-deposition intervals from 949 trail deposition events are compared with theoretical quantiles from an exponential distribution fitted with $\lambda = 1/\langle T \rangle = 0.09$. The mild upward deviation in the upper tail indicates an excess of long intervals relative to an ideal homogeneous Poisson process.

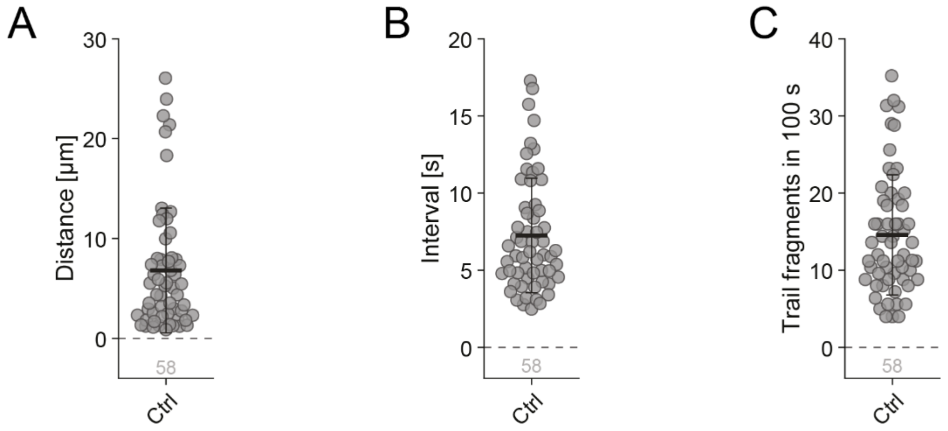


**Fig. S5.**
Trail characteristic measurements pooled from the 58 analyzed sporozoites in Figure 2 F-H. The mean **(A)** distance and **(B)** interval of all trail events of each sporozoites as well as the extrapolated number of **(C)** trail events in 100 s are shown. For the latter, the amount of trail deposition events over the whole imaging time are normalized to 100 s. Shown is the mean value of each characteristic, the standard deviation and the total number n of analyzed sporozoites.

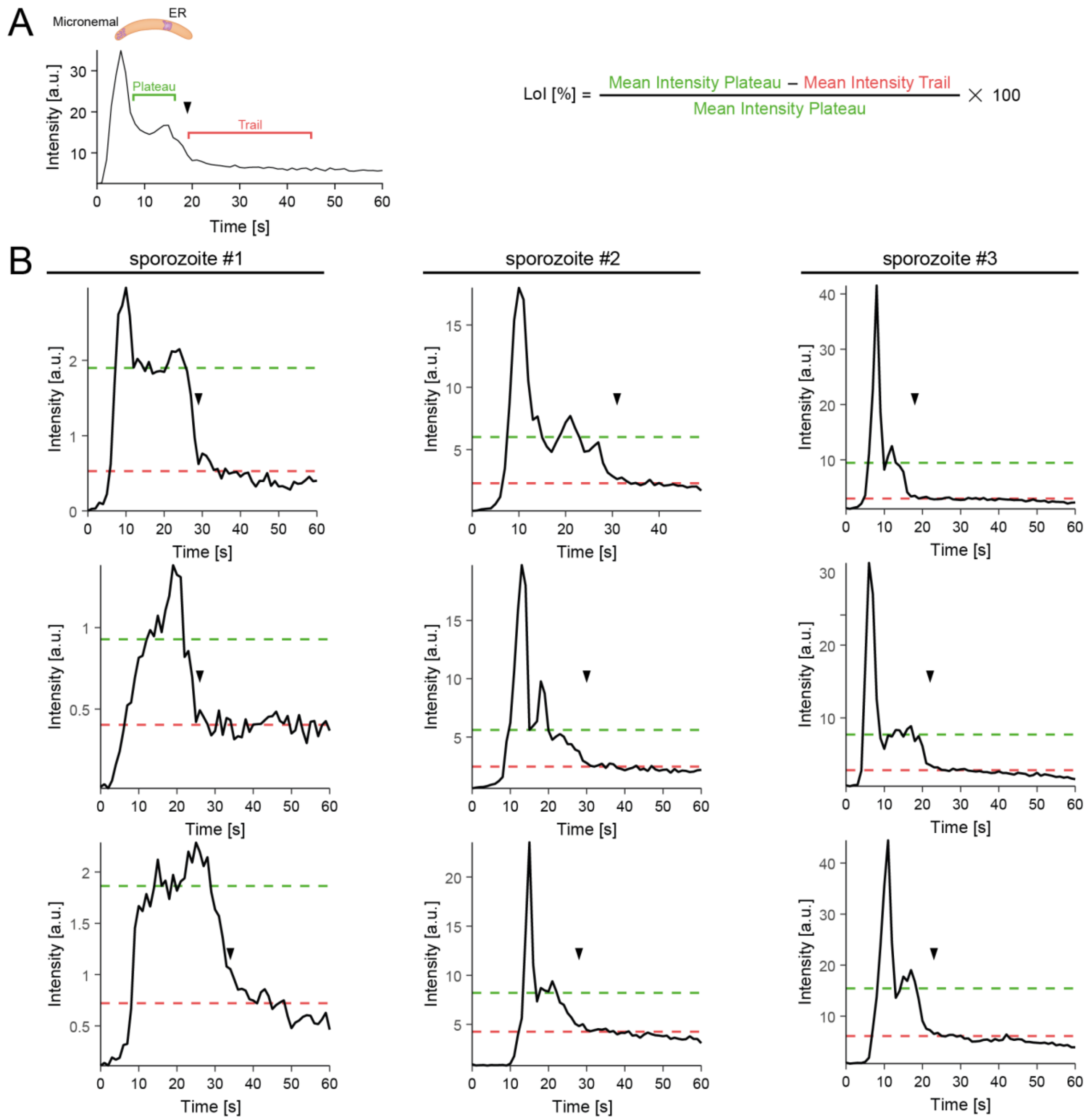


**Fig. S6.**
**(A)** Exemplary intensity profile of a GFP-TRAP analyzed adhesion site with the intensity plateau (green), the remaining signal in the trail (red) and the moment of deposition (black triangle) marked. General formula for the loss of fluorescence intensity (LoI) calculated in the exemplary intensity profile from Figure 4 D. **(B)** Three exemplary adhesion site intensity profiles of 3 individual sporozoites each used for the analysis of the loss of intensity in Figure 5 H. The mean plateau intensity (green dashed line) and the mean trail intensity (red dashed line) used for the loss of intensity calculation are marked alongside the moment of deposition (black triangle).

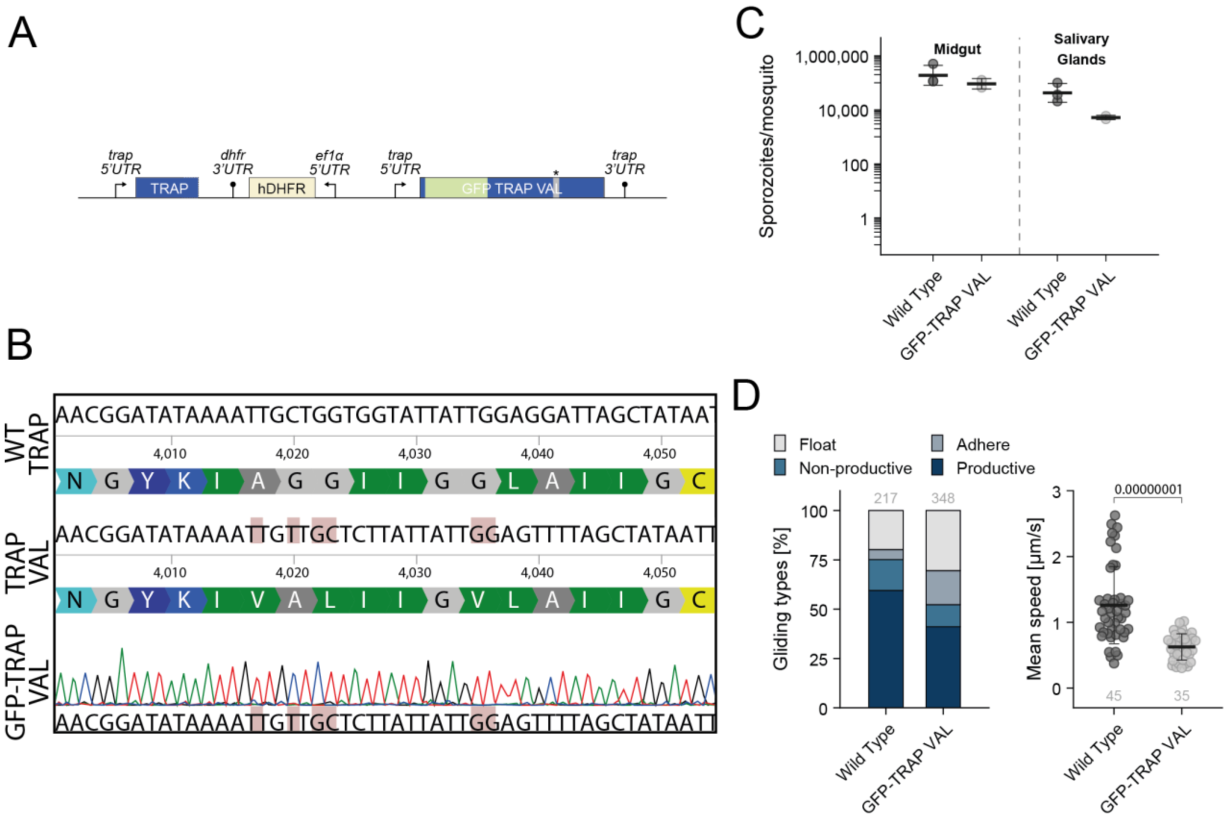


**Fig. S7.**
**(A)** Structure of the transgenic TRAP locus in the GFP-TRAP VAL parasite line. The endogenous TRAP is disturbed through the integration of the construct sequence, integrating a selection marker consisting of the human dihydrofolate reductase (hDHFR) under the ef1α 5' UTR and the *P. berghei* dhfr 3'UTR and the GFP-TRAP VAL sequence under a 5' trap promotor and the endogenous 3'trap UTR. **(B)** Correct integration, presence of the mutations and clonality of the utilized line was confirmed via sequencing. **(C)** The number of sporozoites in the midgut, hemolymph and salivary glands of GFP-TRAP VAL infected mosquitos. The mean values per infected mosquito are shown for the numbers determined at day 18 post infection of two individual cages. **(D)** Gliding motility determined for GFP-TRAP VAL salivary glands sporozoites and categorized according to gliding type. As well as mean speed of productively gliding sporozoite. Data consists of two pooled separate replicates. Significance is calculated through pair-wise comparison with Wild Type in Student's t-test.

**Movie 1 – 3:**
Time series of gliding sporozoites in Figure 2A, B and C. Scale bar = 5 µm. Frame rate = 10 fps.

**Movie 4 – 5:**
Time series of gliding sporozoites in Figure 2D and E. Scale bar = 5 µm. Frame rate = 20 fps.

**Movie 6:**
Time series of gliding sporozoite in Figure 4C. Scale bar = 5 µm. Frame rate = 10 fps.

**Movie 7:**
Time series of gliding sporozoite in Figure 4C. Scale bar = 5 µm. Frame rate = 10 fps.

**Movie 8:**
Time series of gliding sporozoite in Figure 4E and F. Scale bar = 5 µm. Frame rate = 3 fps.

**Movie 9:**
Time series of gliding sporozoite in Figure 4G and H. Scale bar = 5 µm. Frame rate = 10 fps.

**Movie 10:**
Time series of gliding sporozoite in Figure 5B. Scale bar = 5 µm. Frame rate = 10 fps.

**Movie 11 – 12:**
Time series of gliding sporozoites in Figure 5E. Scale bar = 5 µm. Frame rate = 20 fps.

**Movie 13:**
Time series of gliding sporozoite in Figure 5F. Scale bar = 5 µm. Frame rate = 20 fps.

**Movie 14:**
Time series of gliding sporozoite in Figure 6A. Scale bar = 5 µm. Frame rate = 10 fps. Figure time series begins at 48 s.

**Movie 15:**
Time series of gliding sporozoite in Figure 6B. Scale bar = 5 µm. Frame rate = 10 fps. Figure time series begins at 3 s.

**Movie 16:**
Time series of gliding sporozoite in Figure 7A. Scale bar = 5 µm. Frame rate = 5 fps.

**Movie 17:**
Time series of gliding sporozoite in Figure 7A. Scale bar = 5 µm. Frame rate = 5 fps.

**Movie 18:**
Time series of gliding sporozoite in Figure 7B. Scale bar = 5 µm. Frame rate = 5 fps.

**Movie 19:**
Time series of gliding sporozoite in Figure 7E. Scale bar = 5 µm. Frame rate = 10 fps.
